# REnformer, a single-cell ATAC-seq predicting model to investigate open chromatin sites

**DOI:** 10.1101/2025.06.04.657786

**Authors:** Simone G. Riva, Edward Sanders, Tom Wilson, Nicolò Stranieri, E. Ravza Gür, Matthew Baxter, Jim R. Hughes

## Abstract

Genome regulatory elements are fundamental to cellular identity and cell type specific gene expression. Understanding how the underlying genetic code is differentially utilised by different cell types is central to understanding human health and disease. To better understand how DNA encodes genome regulatory elements such as promoters, enhancers, and boundary elements, we leverage the Enformer gene expression and epigenetic prediction model. We used transfer learning with high quality single cell ATAC datasets to develop REnformer, a model to predict chromatin accessibility. By introducing a benchmark for comparing performances against Enformer model, REnformer significantly outperformed Enformer in terms of higher prediction outcomes and lower error rates in all extensive analyses shown; introducing these benchmarks allowed us, and possible future works, to fairly compare such models. We further tested REnformer by predicting the effects of a well characterised *α*-thalassemia variant and found that the prediction aligned with the observed change in genome regulatory element, previously validated. We conclude that REnformer is and can be a state-of-the-art tool to predict cell type specific regulatory elements, and interrogate the effect of genome variation in health and disease.

## I. Introduction & motivation

Chromatin states from DNA sequences provide a more comprehensive view of genome regulation. At the genome level, many non-coding genetic variants associated with human disease and traits might or might not affect this regulation, creating a huge machinery that is manually difficult to decipher and even time-consuming due to the specificity occurring differently in cell populations or tissues. In addition, this variety of mechanisms can be linked to variants or causal associations due to linkage disequilibrium.

To date, numerous computational approaches have been developed to deep-dive into such a problem. Spanning from deep convolutional neural networks, like in DeepSEA [1], Basset [2], and DeepHaem [3], where peak calling is needed to focus on open chromatin regions, to more cutting-edge methods such as transformer-based architecture, like DNABERT [4] and Enformer [5]. The former follows the same paradigm as BERT, a transformer-based model that excels in natural language processing tasks by utilising pre-training and finetuning to develop general understandings from unlabeled data; whereas the latter is a neural network architecture based on self-attention with the aim to predict gene expression and chromatin states from DNA sequences.

With all these new methods and technologies in our hands, it is difficult to understand which approach is correct to answer niche biological questions. Our interest is to better understand CTCF, enhancer, and promoter interactions and their effects on gene expression [6], [7]. Therefore, our work focuses on leveraging the Enformer model to make a better predictor of cell populations and decipher gene regulation in non-coding regions when driven by CTCF, promoter, and enhancer interactions.

A key motivation for this work is its future applications for genome-wide association study (GWAS) and variant prioritisation. As shown in [5], variant effects can be identified in the genome. In GWAS analyses, a haplotype gets linked to a trait. This model enables one to fine-map to both individual variants, cell types, and regulatory elements. Furthermore, if a model correctly links enhancers to promoters, it can enable one to link GWAS haplotypes to genes (via their promoters). In this study, we present REnformer, a network based on a previously published model (Enformer) [5], with the intent to predict open chromatin regions from DNA sequence from 151 single-cell Assay for Transposase-Accessible Chromatin using sequencing (scATAC-seq); introducing a data normalisation step and a benchmark that shows the better performance that can be achieved with REnformer, but also allowing bench-marking future approaches with the current state-of-the-art.

## II. Data & Method

This section focuses on what publicly available data we collected, how we processed it, and the architecture of REnformer.

### A. Data collection

We collected 151 different scATAC-seq types from various samples and collections (genome version GRCh38), allowing us to cover almost the entire human body. In detail, we selected 7 cell types from Turner *et al*. [8], 107 cell types from Zhang *et al*. [9], 15 cell types from Granja *et al*. [10], 18 primary peripheral blood mononuclear cells (PBMC) cell types (See Table I), and 3 erythroid differentiation stages and 1 human umbilical vein endothelial cell (HUVEC) *in-house* generated (see Table II) for full list of cell types).

**TABLE I.**
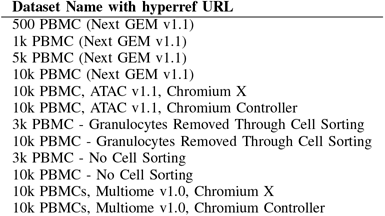
Human Healthy Donors, 10x Genomics, accessed 04/03/2023.

**TABLE II.** LIST OF SCATAC-SEQ CELL TYPES USED IN Renformer.

| Cell type | Cell type | Cell type | Cell type | Cell type |
| --- | --- | --- | --- | --- |
| Endothelial Myocardial | Plasma | Fetal Endothelial General | Fetal Skeletal Myocyte | pDC |
| Enterocyte | Schwann General | Fetal Endothelial Hepatic | Fetal Stellate | Pre B |
| Esophageal Epithelial | SI Goblet | Fetal Endothelial Placental | Fetal Sympathoblast | B intermediate |
| Fasciculata | Smooth Muscle Colon | Fetal Enteric Glia | Fetal Syncitio and Cytotrophoblast | B memory |
| Fibro Epithelial | Smooth Muscle GE junction | Fetal Enteric Neuron | Fetal T Lymphocyte CD4+ | B naive |
| Fibro General | Smooth Muscle GI | Fetal Enterocyte | Fetal T Lymphocyte Cytotoxic | CD14 Monocytes |
| Fibro GI | Smooth Muscle Muscularis | Fetal Erythroblast | Fetal T Lymphocyte IL2+ | CD16 Monocytes |
| Fibro Liver Adrenal | T Lymphocyte CD4+ | Fetal Excitatory Neuron | Fetal Thymocyte | CD4 Naive |
| Fibro Muscle | T Lymphocyte CD8+ | Fetal Fibro General | Fetal Ureteric Bud | CD4 TCM |
| Fibro Nerve | Transitional Cortical | Fetal Fibro GI | Fetal V Cardiomyocyte | CD4 TEM |
| Follicular | Type I Skeletal Myocyte | Fetal Fibro Muscle | Erythroid day0 | CD8 Naive |
| Hepatocyte | Type II Skeletal Myocyte | Fetal Fibro Placental | Erythroid day10 | CD8 TCM |
| Keratinocyte | V Cardiomyocyte | Fetal Fibro Splenic | Erythroid day5 | CD8 TEM |
| Luteal | Vascular Smooth Muscle | Fetal Gastri Goblet | B | cDC2 |
| Lymphatic | Fetal A Cardiomyocyte | Fetal Hepatoblast | CD4 M | HSPC |
| Macrophage General | Fetal Acinar | Fetal Inhibitory Neuron | CD8 CM | MAIT |
| Mammary Basal Epithelial | Fetal Adrenal Cortical | Fetal Macrophage Hepatic | CD8 EM | NK |
| Mammary Epithelial | Fetal Adrenal Neuron | Fetal Macrophage Placental | cDC1 | Plasmablast |
| Mammary Luminal Epithelial | Fetal Alveolar Cap Endo | Fetal Megakaryocyte | CLP | Platelet |
| Memory B | Fetal Alveolar Epithelial | Fetal Mesangial | CMP LMPP | Treg |
| Mesothelial | Fetal Astrocyte | Fetal Metanephric | Early Basophil | HUVEC |
| Naive T | Fetal Cardiac Fibroblast | Fetal Photoreceptor | Early Erythroid |  |
| Natural Killer T | Fetal Chromaffin | Fetal Retinal Neuron | GMP |  |
| Nerve Stromal | Fetal Ciliated | Fetal Retinal Progenitor | GMP Neutrophils |  |
| Parietal | Fetal Ductal | Fetal Satellite | HSC |  |
| Pericyte Muscularis | Fetal Endocardial | Fetal Schwann General | Late Erythroid |  |

### B. Data processing

The *in-house* generated data has been processed using 10X Genomics Cell Ranger ATAC/ARC (v2.0.0), making bigWig files; Turner *et al*., Granja *et al*., and PBMCs datasets have been (upstream and downstream) analysed in more detail to isolate and generate bigWig files of the correct cell types of interest; whereas, Zhang *et al*. data has been taken from their interactive web atlas in bigWig format directly.

Once all bigWig files (one per cell type) were collected, we normalised the coverage counts across all samples to make them comparable across experiments and cell types using the *in-house* zone z-score normalisation (*ZZS*). This is a modified version of z-score normalisation used to make RNA-seq comparable between samples [11]. Compared to the traditional z-score, the modified version is better suited for skew and more robust to outliers [Reference book: Volume 16: How to Detect and Handle Outliers, Boris Iglewicz, David Caster Hoaglin].

*ZZS* is divided into 6 phases: 1) Coordinates within the hg38 blacklist [12] and regions with 500 base-pairs (bp) or more consecutive zeros were masked. To reduce the impact of noise, remaining signal was smoothed via convolution with a 301 bp triangular kernel. 2) The modified z-score fits a normal distribution per chromosome *c* centred at the median of *c* and scaled by an approximation of 1 standard deviation. The median absolute deviation from the median for *c* was calculated as 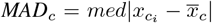 and if *MAD* ≠ 0, the scale set as 1.4826 × *MAD*. Otherwise, the mean absolute deviation was calculated as 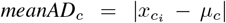 and scale set as 1.2533 × *meanAD*. These constants were taken from common usage of scale parameters [13]. 3) As the vast majority of scATAC-seq signal is noise, a threshold was derived using a probability of 99.7% to capture the value at which the smoothed signal for *c* falls three deviations from the median. The threshold was applied across each bp in the smoothed signal. Start and end coordinates exceeding the threshold for 35 bp or more were extracted, rounded to the nearest full-sized bin of 1, 000 bp size and consecutive coordinates merged forming signal zones. 4) Fragment size *f*_*c*_ was estimated per chromosome as the smallest non-zero value 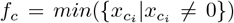 and the global mean reads excluding zeros *µ*_*g*_ calculated across all chromosomes. Zones were then filtered to remove those where all signals falls below quality threshold *τ*_*c*_ = *max*(1.5*µ*_*c*_, 1.5*µ*_*g*_, 10*f*_*c*_) — we observed the upper noise boundary tends to fall around 1.5× the non-zero mean, so 1.5 × is a safety net for larger library sizes; and anything less than 10 reads was considered low-quality data, as also shown in pre-filtering count cut-off recommended by DESeq2 [14] and csaw [15]. 5) For each sample, the bigWig signal across its zones is extracted per chromosome and zeros removed. Then, the 10^*th*^ lower and upper percentile of values are also removed to improve robustness against technical artifacts. An intermediate signal per bp is calculated via the z-score:

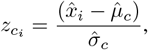

by using the remaining signal 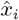, its mean 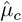 and standard deviation 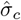) Finally, as scATAC-seq signal is positive, if *z*_*c*_ contains a negative value, the minimum is subtracted to produce the normalised chromosome signal *ZZS*_*c*_:

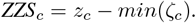

We then processed all normalised samples and chunk the sequences in 196,608 bp, covering 50% of the genome per sample, removing blacklist regions, and obtaining a total of 7,099 regions functioning as input for the model. These 7,099 regions have also been converted into 896 bp in length, corresponding to 114,688 bp, by cropping 40,960 bp on each side and aggregated into 128 bp bins as the target of the model.

### C. Method

We build REnformer leveraging Enformer architecture, which consists of 7 convolutional blocks with pooling, 11 transformer blocks, and a cropping layer followed by final pointwise convolution developed in Tensorflow (v2.4.1) [16] and Sonnet (v2.0.0) [17] and converted into PyTorch (v2.4.1) [18] under Python v3.11.11. We also mapped the trained Tensorflow weights, released by Avsec *et al*. [5], into PyTorch weights for transfer learning purposes and replaced the output head of Enformer with our output head (see Figure 1).

**Figure 1.**
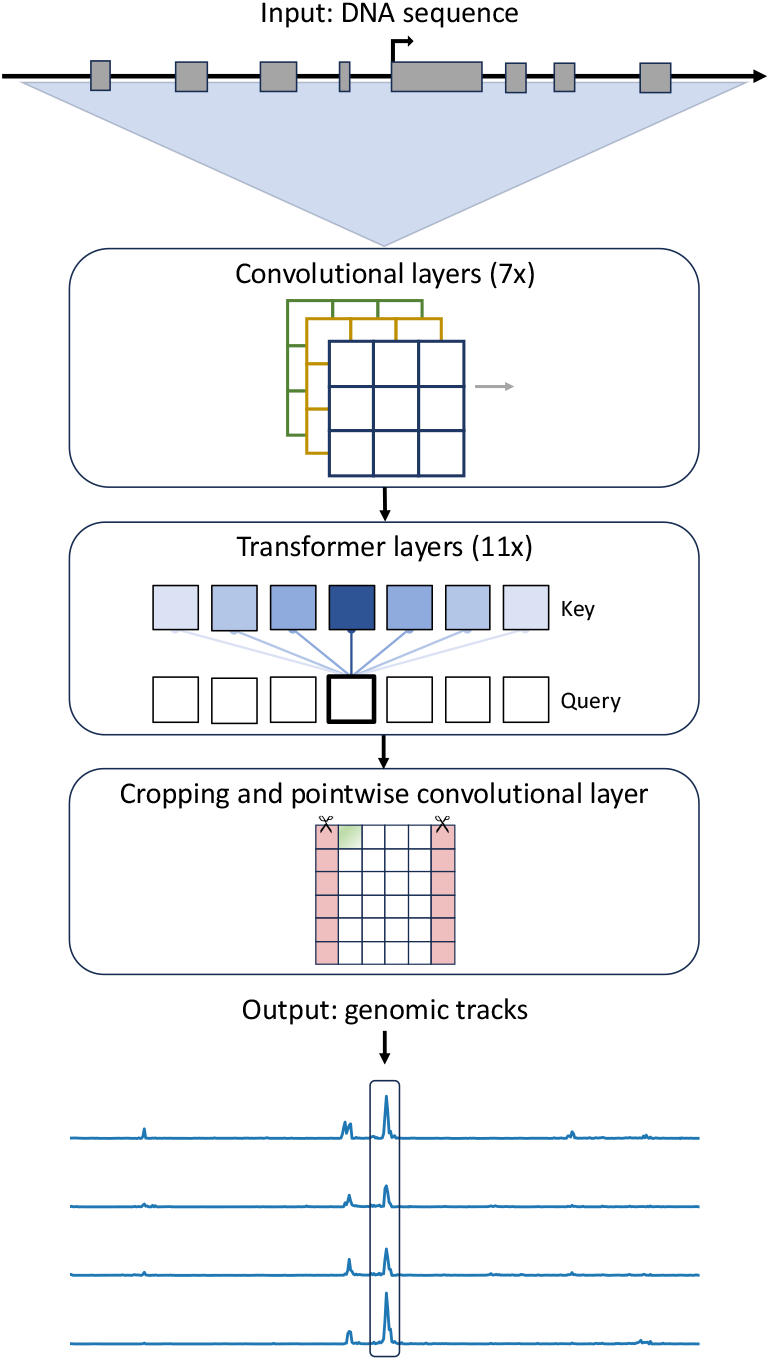
Overview of REnformer model.

With a one-hot-encoded DNA sequence of length 196,608 bp as input (A = [1,0,0,0], C = [0,1,0,0], G = [0,0,1,0], T = [0,0,0,1], and N = [0,0,0,0]), our model predicts 151 genomic tracks for the human genome, each of the tracks being 896 bp long, totalling 114,688 bp aggregated into 128 bp bins, as explained in Section II-A and II-B.

## III. Result

We introduce REnformer, a revisited version of Enformer, leveraging transfer learning strategy to train the network on scATAC-seq data only — since it offers more thorough insight into the use of such data and is abundant in epigenetic information at the cell level, when compared to any bulk technology [19] — with the purpose of predicting open chromatin from DNA sequence in humans and investigating possible hypotheses on genomic variation within chromatin states. In the training phase, we applied the transfer-learning technique, consisting of a subset of fine-tuning approach class, only the new final layers are trained on the new scATAC-seq data, whilst the rest of the model has been frozen, leveraging the learned features from the original Enformer model. We sampled the genome, selecting 9, 406 regions and divided them into train and test sets, respectively, 7, 099 (75%) and 2, 307 (25%) 196,608 bp region length. We then trained REnformer on Nvidia L40S graphics processing unit using the 7, 099 regions. Figure 2 shows the output of Renfomer in chr10 : 22241840 *−* 22356528 region — selected from the test set — comparing the output of HUVEC experiment provided to REnformer (light green) and its prediction (dark green), against the two DNase I hypersensitive sites sequencing (DNase) HUVEC provided to Enformer (DNase experiment HUVEC 1 in light blue and Enformer prediction in dark blue; DNase experiment HUVEC 2 in light orange and Enformer prediction in dark orange). As illustrated in Figure 2, using the normalisation described in Section II-B, and scATAC-seq data for REnformer, allowed us to fine-map the open chromatin sites with a better global visual resolution than Enformer. Even if Enformer can predict open chromatin, there is a lack of sensitivity in some hypersensitive regions.

**Figure 2.**
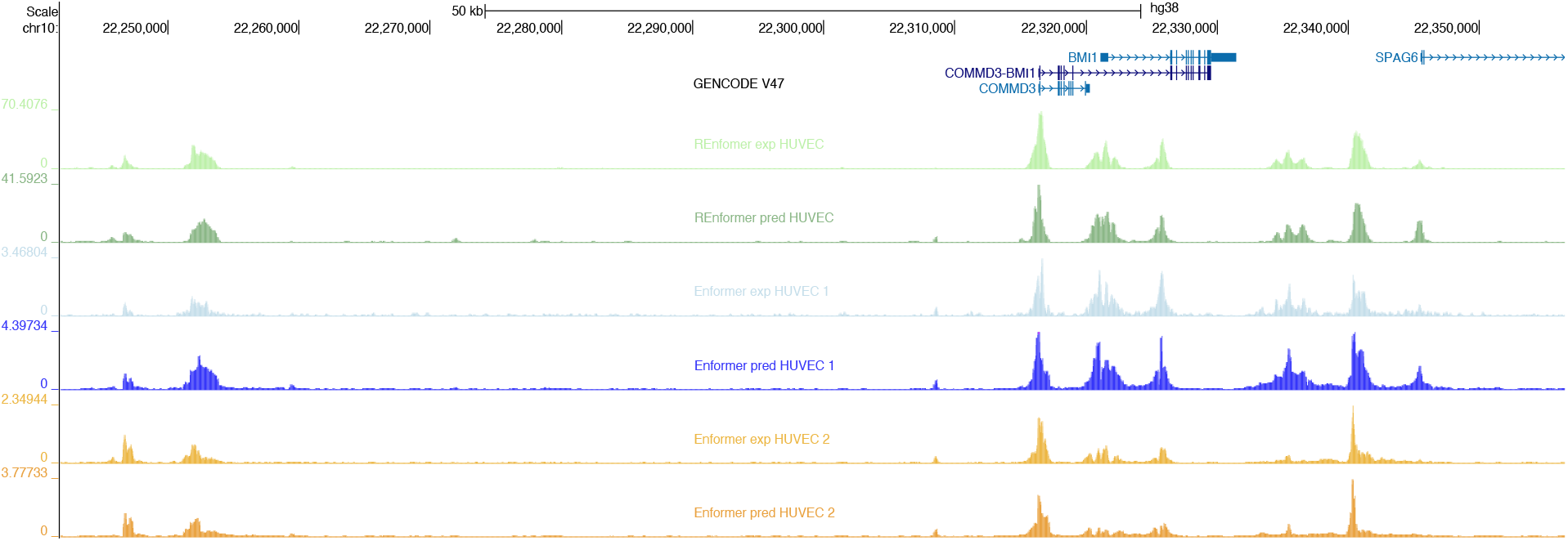
Genome Browser track for chr10 : 22241840 *−* 22356528 region. In green, experiment (light) and predicted (dark) HUVEC cell type for REnformer model. In blue and orange, experiments (light) and predicted (dark) 2 DNase HUVEC cell types for Enformer model.

Subsequently, we introduced three tests to evaluate and better compare REnformer against Enformer performances, all based on the root mean square error (*RMSE*):

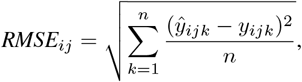

where *ŷ*_*ij*1_, …, *ŷ*_*ijk*_, …, *ŷ*_*ijn*_ represent the predicted values, and *y*_*ij*1_, … *y*_*ijk*_, …, *y*_*ijn*_ denote the ground truth values for element *i* in the set, class *j* ∈ [1, …, 151], and region *k* ∈ [1, …, *n*] with *n* = 896. All three metrics are a variation of the normalised *RMSE*_*ij*_ (*NRMSE*_*ij*_): firstly, normalising by using the standard deviation *σ*:

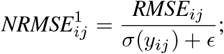

secondly, normalising by the min-max values:

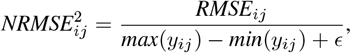

and thirdly, normalising by the difference of the 75^*th*^ and 25^*th*^ quantiles:

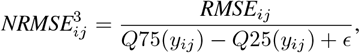

where *ϵ* = 1*e*^*−*8^ in all three cases, since some values in the ground truth are constant in some region *i*, and to avoid division by zero error. From the test set, we removed regions where the target was only filled with zeros (123), obtaining 2, 184 regions and randomly grouped them, without duplication in six subsets. We then applied the 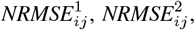, and 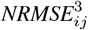 between all predicted *ŷ*_*ij*_ and observed *y*_*ij*_ values, where *i* is the region and *j* the class. Similarly, we applied the same tests on 151 (same number of classifiers in REnformer) selected DNase classifiers in Enformer model and visualised the results in Figure 3. For each 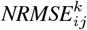, where *k* ∈[1, 2, 3], we also applied Mann-Whitney U test [20], by using `alternative = ‘less` ‘. The distribution underlying REnformer observed and predicted data is stochastically less than the distribution underlying Enformer observed and predicted data, for all regions of interest, using Bonferroni correction [21], showing that in all comparisons— except in subset 6 for 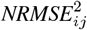, where we cannot rule out the null hypothesis — there is a statistically difference in 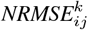 in favour of REnformer outcomes. This analysis introduced a way to compare and benchmark REnformer against Enformer, or any future model based on either of these, and showed that applying transfer learning along with the introduced data processing (explained in Section II-B), REnformer outperformed Enformer model. Furthermore, the kernel density estimates (*KDE*) in Figure 4 indicated that the *KDE* curves for REnformer begin at lower error values, reach higher peaks, and drop off more rapidly, indicating overall better performance. However, the greater variability in the REnformer peaks suggests a higher heterogeneity in the difficulty of predicting values among classes.

**Figure 3.**
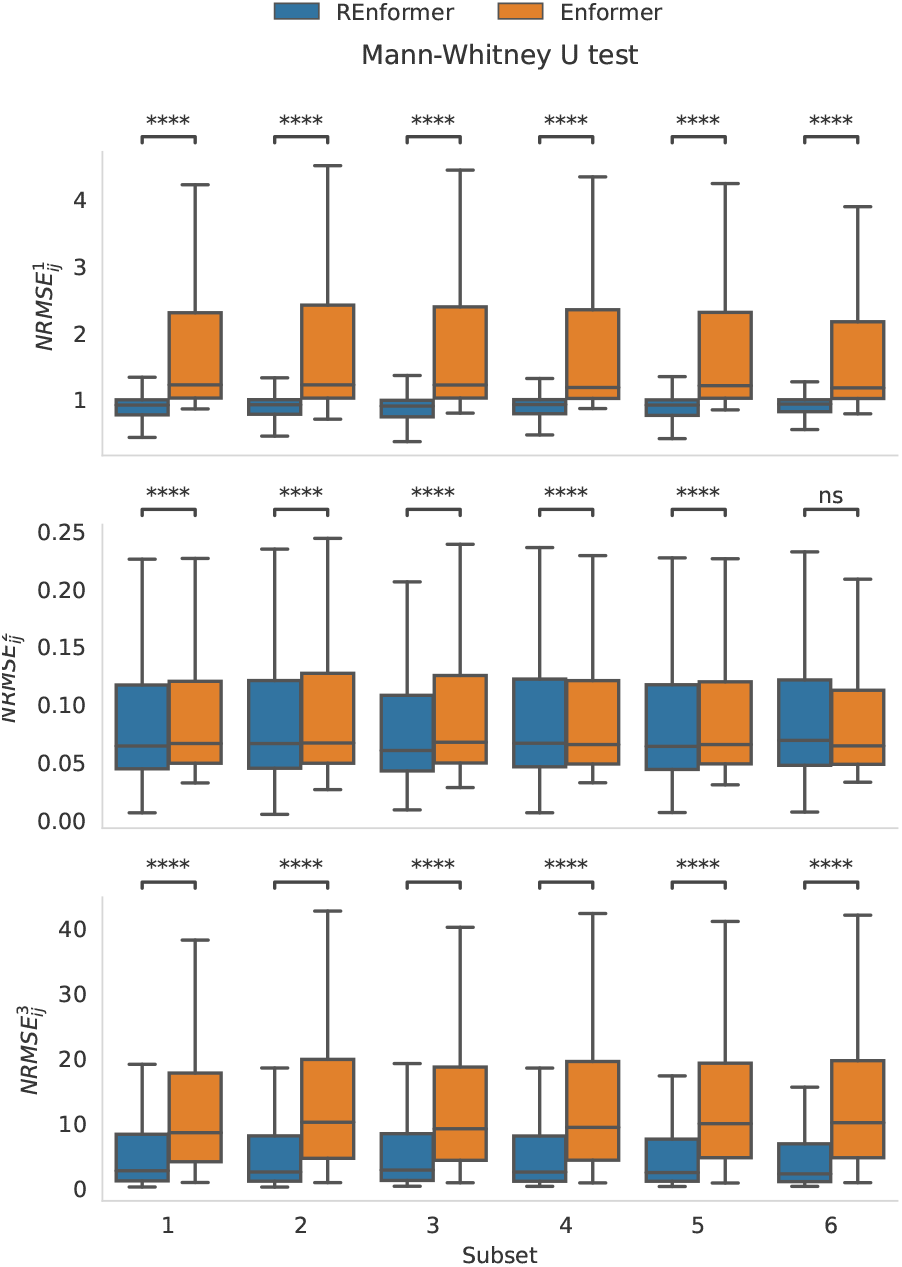
Boxplots representing the 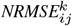, where *k* ∈ [1, 2, 3] for each random test subset; showing the performance of REnformer (blue) against Enformer (orange). On top of every 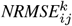 and subset is the p-value of Mann-Whitney U test, where ∗means the p-value ≤ 1*e*^*−*4^ and ns means p-value *>* 0.05.

**Figure 4.**
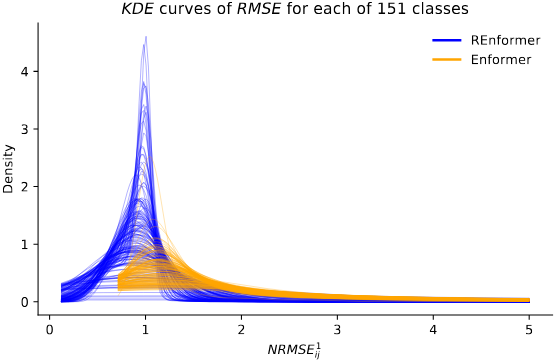
*KDE* of the 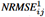 reveal that REnformer (blue) exhibits lower minimum and mean error values than Enformer (orange).

In addition, we checked edge cases results to understand the differences between REnformer predictions and the observed values. We plotted selected the randomly selected regions in subset 1, and in Figure 5, where each dot represents a region *i* for a classifier *j* for REnformer model, with the *RMSE*_*ij*_ (*x*-axis) against the 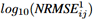 (*y*-axis. We picked 5 dots *ij*, a random one (orange), the one with the worst *RMSE*_*ij*_ value (green), the worst 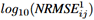 (red), the best 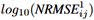 (purple), and the best *RMSE*_*ij*_ (brown). Consequently, we showed the predicted and observed genomic tracks for these 5 regions (see Figure 6), instructing us that such difference between the prediction and the ground truth is minimum; for the worst *RMSE* (6**a**), worst 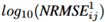 (6**b**), random (6**c**), and the best 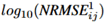 regions the difference is only in the magnitude of the open chromatin sites. In contrast, the best *RMSE*_*ij*_ (6**e**) region is related to a genomic region filled with zeros in the target; whilst its prediction, for the selected class, is asymptotically zero in all the prediction scores across the targeted genomic region. This shows that also in the edge-cases, the prediction is close to the observed values, confirming the positive outcomes of our REnformer model.

**Figure 5.**
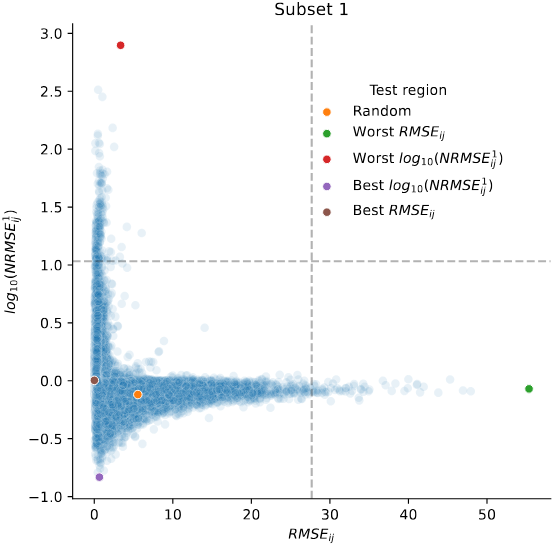
Scatterplot showing on the x-axis *RMSE*_*ij*_ and on the y-axis the 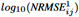, whilst dots are representing regions *i* and classifiers *j* of REnformer in the random test subset 1. Highlighted in orange, green, red, purple, and brown a random case and 4 edge cases or the *RMSE*_*ij*_ and 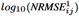.

**Figure 6.**
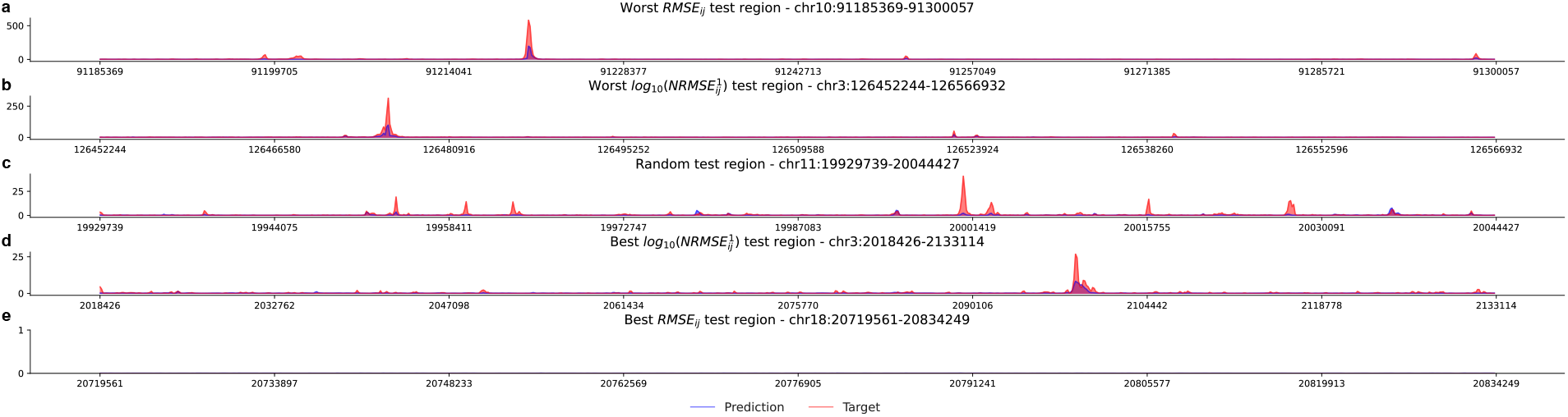
Lineplots of the highlighted regions in Figure 5. Blue lines represent the predicted values, whilst the red lines represent the observed values. **a)** shows the worst 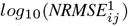 (region chr10 : 91185369 − 91300057); **b)** the worst 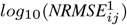 (region chr3 : 126452244 − 126566932); **c)** the random region (region chr11 : 19929739 −20044427); whereas **d)**, the best 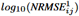 (region chr3 : 2018426 −2133114). In these 4 cases, the difference between predictions and ground truths corresponds to a magnitude difference. Instead, **e)**, which represents the best *RMSE*_*ij*_ (region chr18 : 20719561 − 20834249) corresponds to a region where no open chromatin sites are present, either for the observed and the predicted values.

Given the excellent obtained REnformer results, we wanted to investigate some mutagenesis analysis and how variants might affect open chromatin sites. To do so, we choose a variant form of the inherited blood disorder *α* thalassemia [22], since our model has learnt three behaviours of the erythroid differentiation. This selection has been driven from the fact that this variant T into C, occurring in position chr16 : 159710 (in hg38) is well understood and studied in the biological field, as it is a gain of function, which act as an orientation dependent enhancer blocker, by creating a new promoter [22], [23]. With this information, providing the reference genome and the alteration genome (which includes the variant T instead of the C), we expected to see a peak underneath the variant region only when predicting the open chromatin in erythroid data in the altered genome. Indeed, Figure 7 shows T/C mutation, where it is visualised, in green, the most differentiated experiment erythroid track (chr16 : 102366 *−* 217054), the prediction using REnformer using the hg38 reference genome in blue, showing that it resembles coherently the epigenetic sites as per the experiment, and the prediction with the variation in it in red. To visualise better where the gain of function and the promoter appear, we subtracted from the prediction on the alteration, the prediction on the reference, in purple, showing that the created promoter, in fact, becomes visible where the T/C variation is applied; even more strongly visible when overalying the blue and red, respectively the reference and altered genome, tracks. This result, indeed, was aligned with the discovery in early 2000s by De Gobbi *et al*. [22] and then reconfirmed in more details by Bozhilov *et al*. [23].

**Figure 7.**
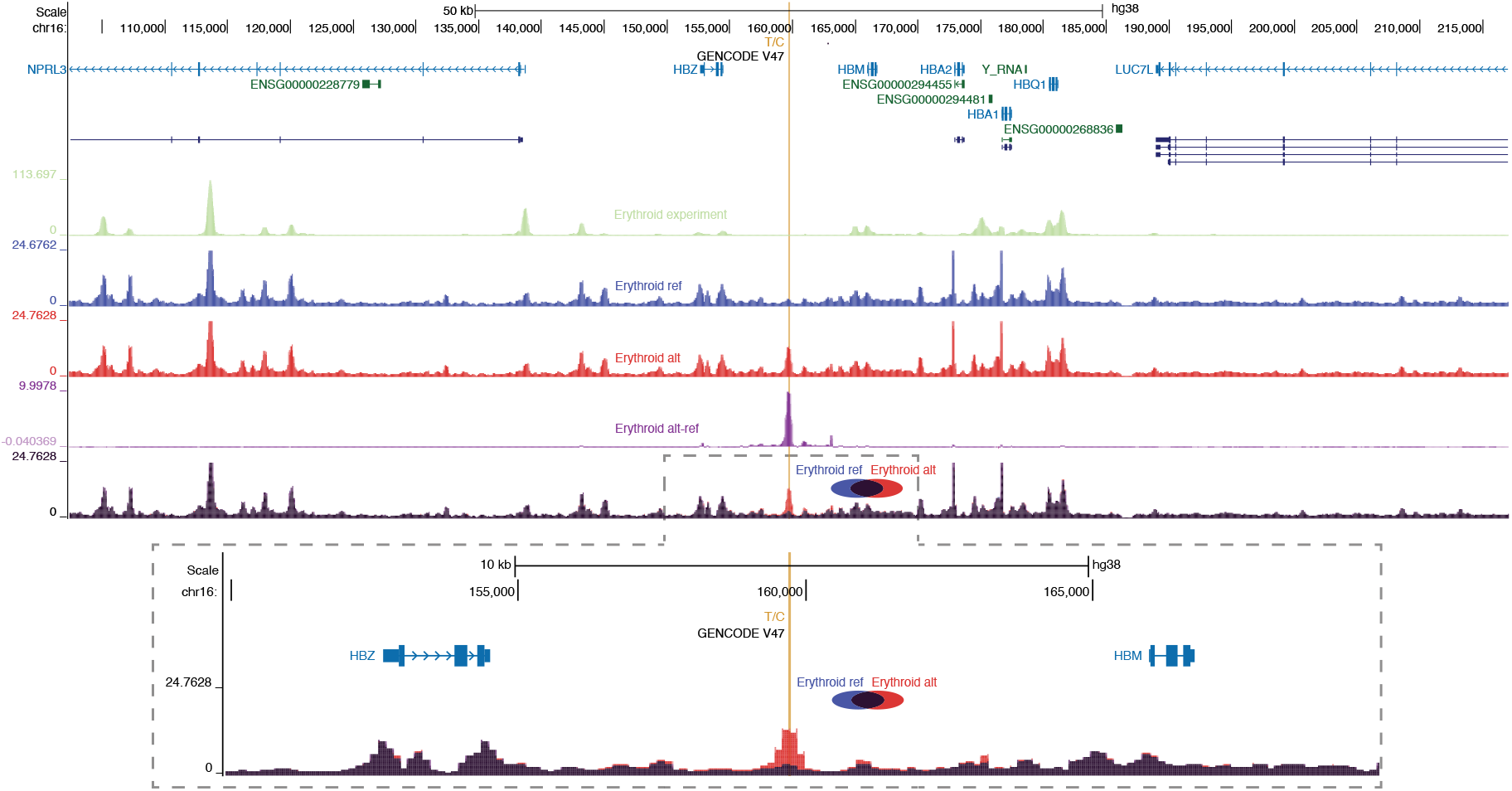
Genome Brower *α* thalassemia locus (chr16 : 102366− 217054). This figure shows the effect of the variation from T to C in position 159710 (highlighted in yellow) in erythroid cells. As a convincing factor, the first track, in green, represents the erythroid experiment data, the prediction on the reference genome in blue (where the T is located), the prediction on the variation in red (where the T has been replaced with the C). We then subtracted the blue with the red tracks, showing where the changes occurred (in purple), and below, the overlay of the blue and red predictions.

## IV. CONCLUSION & FUTURE WORK

In this work, we presented REnformer, a transformer-based approach, based on the published Enformer model, where applying transfer learning strategy to learn epigenetic features from scATAC-seq data, we were able to achieve a better performance (Figure 3) and resolution Figures 5 and 6 on predicted values (Figure 2), given the introduced data normalisation (Section II-B) and the high resolution of the data collected (Section II-A). Furthermore, we showed that single nucleotide variant (SNP) analysis, i.e., looking at the *α* thalassaemia example, where the T into C, shown in Figure 7, demonstrating that REnformer can detect this change, which creates a promoter. In fact, a key application is to comprehend mutagenesis, the effect of variants, to better fine-map GWAS traits, with also applications to rare diseases. Linking these achievements with REgulamentary [24], where using open chromatin and histone mark data can allow us to link haplotypes to gain or loss in gene regulatory regions with high granularity. This is often limited by the inability to fine-map from a linked haplotype, containing many variants, to one or more variants. Currently, we have to integrate REnformer with other pipeline analyses to achieve these goals. REnformer will enable us to prioritise and rank which variants, within a haplotype, are most likely to have a causal relationship, and eventually also to infer the interaction’s directionality between regulatory elements and decipher better the non-coding landscape. Lastly, to further improve and refine REnformer prediction of open chromatin in different classes and minimise errors between predicted and observed values, an adjustment on the proposed loss function, by Avsec *et al*., and batch size during the training phase should be taken into consideration.

## Acknowledgment

S.G.R. is supported by the MRC grant (MC UU 00029/3). E.S. and M.B. are supported by the Wellcome Trust grant (225220/Z/22/Z). T.W is supported by the MRC grant (MC ST 00029). E.R.G. is supported by the Ministry of National Education Selection and Placement of Candidates Sent Abroad for Postgraduate Education (YLSY) scholarship, Republic of Türkiye Ministry of National Education and the Wellcome Trust grant (225220/Z/22/Z). J.R.H. is supported by the Wellcome Trust grants (225220/Z/22/Z and 106130/Z/14/Z) and the MRC grant (MC UU 00029/3).

## Declaration

J.R.H. is a co-founder and director of Nucleome Therapeutics and provides consultancy to the company.

## Dataset Name with hyperref URL

